# Recombinant protein production and purification of SiiD, SiiE and SiiF - components of the SPI4-encoded type I secretion system from *Salmonella* Typhimurium

**DOI:** 10.1101/2020.01.25.919720

**Authors:** Stefan Klingl, Sina Kordes, Benedikt Schmid, Roman G. Gerlach, Michael Hensel, Yves A. Muller

**Affiliations:** Division of Biotechnology, Department of Biology, Friedrich-Alexander-University Erlangen-Nürnberg, D-91052 Erlangen, Germany; Robert Koch-Institut, Wernigerode, Germany; Abt. Mikrobiologie and CellNanOs, Universität Osnabrück, Osnabrück, Germany

**Keywords:** *Salmonella*, type I secretion system (T1SS), Protein expression, Purification

## Abstract

In humans, *Salmonella enterica* infections are responsible for a pleiotropy of medical conditions. These include intestinal inflammation and typhoid fever. The initial contact between *Salmonella* and polarized epithelial cells is established by the SPI4-encoded type I secretion system (T1SS), which secrets SiiE, a giant non-fimbrial adhesin. We have recombinantly produced various domains of this T1SS from *Salmonella enterica* serovar Typhimurium in *Escherichia coli* for further experimental characterization. We purified three variants of SiiD, the periplasmic adapter protein spanning the space between the inner and outer membrane, two variants of the SiiE N-terminal region and the N-terminal domain of the SiiF ATP-binding cassette (ABC) transporter. In all three proteins, at least one variant yielded high amounts of pure soluble protein. Proper folding and cooperative unfolding were investigated by circular dichroism (CD) spectroscopy. Secondary structure content estimations from CD spectra were in good agreement with the values derived from SiiD and SiiF homology models or, in case of the SiiE N-terminal region, a secondary structure prediction. For one SiiD variant, protein crystals could be obtained that diffracted X-rays to approximately 4 Å resolution.

## 1. Introduction

*Salmonella enterica* is a food-borne pathogen that causes diseases such as self-limiting gastroenteritis, intestinal inflammation and typhoid fever. Multiple virulence factors and protein secretion systems, which are mostly encoded by so-called *Salmonella* Pathogenicity Islands (SPI), enable *Salmonella* to infect a broad range of hosts [1, 2]. While a SPI1-encoded type III secretion system delivers a cocktail of effector proteins into the host cells and is generally required for invasion, the initial contact to polarized epithelial cells is established *via* a SPI4-encoded type I secretion system (T1SS). T1SSs are widely spread among Gram-negative bacteria. They translocate proteins from the cytoplasm into the extracellular space with the translocated proteins fulfilling tasks like nutrient uptake, host invasion or biofilm formation [3]. Translocation occurs in a one-step process without periplasmic intermediates. The *Salmonella* SPI4 locus harbors a total of six proteins designated SiiA-F [4]. SiiC, SiiD and SiiF constitute the canonical building blocks of the T1SS, SiiE is the secreted substrate and SiiA and SiiB are two accessory proteins that enhance the invasion of polarized cells [4, 5](Supplementary Information S. Fig. 1). Of the three canonical T1SS proteins, SiiF is the inner membrane transport ATPase belonging to the family of ATP-binding cassette (ABC) transporters, SiiD is the periplasmic adapter protein (PAP, also termed ‘membrane fusion protein’) and SiiC is the outer membrane protein (OMP).

Typically, SiiF-like ABC transporters are composed of two transmembrane domains and two C-terminal nucleotide-binding domains. The various domains are responsible for substrate recognition and specificity as well as for energizing the secretion process by ATP hydrolysis [6]. In some instances, the T1SS-specific ABC transporters encompass at their N-terminus an additional domain, namely either a C39 peptidase or a C39 peptidase-like domain (CLD) [7]. ABC transporters with a C39 peptidase domain usually transport antimicrobial peptides, and the peptidase cleaves off the substrate’s leader peptide prior to secretion. In CLD-containing ABC transporters, the CLD engages into specific interactions with the transported substrate and is essential for secretion. For example, in the hemolysin ABC transporter HlyB from *E. coli*, CLD acts as a chaperone and prevents the aggregation and/or degradation of the secreted toxin HlyA [8]. SiiF belongs to the T1SS-specific ABC transporters that contain an additional N-terminal domain. However, it is currently not known whether this domain belongs to the C39-peptidase type, the CLD type or adopts an altogether different 3D structure.

The PAP acts like a clamp between the ABC transporter and the pore-forming OMP. SiiD-like PAPs from T1SSs are structurally homologous to PAPs from bacterial tripartite multidrug efflux pumps. The ABC-subfamily of these efflux pumps resembles the molecular architecture of T1SSs with an ABC transporter, a PAP and an OMP [9]. A recent electron cryo-microscopy structure of the ABC-type tripartite macrolide-specific efflux pump from *E. coli* reveals a hexameric assembly of the PAP MacA that bridges between the dimeric ABC transporter MacB and the trimeric OMP TolC and generates a channel for substrate translocation [10]. SiiD-like PAPs share a common domain architecture. In these proteins, an N-terminal transmembrane helix, which anchors the protein in the inner membrane, is followed by a periplasmic domain consisting of a β-barrel, a lipoyl and a coiled-coil domain. The OMP SiiC in the *Salmonella* T1SS is predicted to be structurally homologous to TolC, which is participating in the *E. coli* HlyB–HlyD–TolC hemolysin system [11].

The only known substrate for the SPI4-T1SS is SiiE, a 595 kDa non-fimbrial adhesin [12]. It is composed of an N-terminal domain, containing β-sheet and coiled-coil structures, followed by 53 bacterial immunoglobulin-like (BIg) domains with a putatively unfolded insertion segment occurring between BIg52 and BIg53. A signal sequence at the C-terminus allows for the secretion by the SPI4-T1SS. SiiE induces the first contact between bacteria and host cells. C-terminal portions of SiiE bind to glycostructures at the apical side of polarized cells containing N-acetyl-glucosamine and/or α2,3-linked sialic acid [13]. A crystal structure of SiiE BIg50-52 provided insight into the BIg domain architecture and revealed two types of Ca^2+^-binding sites, which stabilize the rod-like structure [14]. Further studies showed that type I Ca^2+^-binding sites were essential for the efficient secretion of SiiE, whereas integrity of type II sites in the C-terminal portion was needed for mediating adhesion and invasion [15]. During the phase of highest bacterial invasiveness, SiiE is retained on the bacterial envelope and, at later time points, secreted into the culture medium. Deletions in the N-terminal domain of SiiE affect retention of SiiE [16]. However, the underlying mechanism of the transition from the surface-bound to the soluble form is not well understood.

In this study, we report the successful high yield recombinant production of soluble domains of SiiD, SiiE and SiiF from *Salmonella enterica* serovar Typhimurium (*S.* Typhimurium) for further biochemical and biophysical investigations.

## 2. Materials and Methods

### 2.1. Bioinformatics studies

Structural homologs of SiiD, SiiE and SiiF were identified using the HHpred server [17]. The sequence alignment of SiiD with structurally homologous proteins was performed using program CLUSTAL-OMEGA [18], and the alignment conservation score was extracted from the JALVIEW bioinformatics software [19]. Transmembrane regions in SiiD and its structural homologs were identified with the TMpred server (https://embnet.vital-it.ch/software/TMPRED_form.html). The JPred4 server was used to predict SiiD secondary structure elements [20]. Homology models of SiiD and SiiF were obtained from the SWISS-MODEL server [21]. Program DSSP was used to classify the secondary structure elements of the SiiF model and the PCAT1 3D structure [22]. The GeneSilicoMetaDisorder server was used for the prediction of disordered regions in the N-terminal domain of SiiE [23].

### 2.2. Cloning

The *siiD* gene was PCR-amplified from genomic DNA of *S.* Typhimurium NCTC12023 using primers (endonuclease restriction sites in all primers are underlined) B2H-XbaI-SiiD-For: 5’-GAC<u>TCTAGA</u>GAATAGAAGACAAAGCGATCATC-3’ and B2H-SiiD-KpnI-Rev: 5’-CTTA<u>GGTACC</u>CAAGGTGTATCTAATCGTTTAG-3’. The resulting fragment was digested by XbaI and KpnI restriction enzymes (New England Biolabs) and subsequently ligated in the similarly digested pUT18C [24] vector (pWRG679). SiiD (UniProt accession no. **<u>O85314</u>**) variants SiiD(30-393), SiiD(43-300) and SiiD(80-263) (amino acid (aa) coding region in parentheses, respectively) were obtained from vector pWRG679 by PCR using primers SiiD(30-393)-BamHI-For: 5’-GGAGGAGGAGA<u>GGATCC</u>AATTCAGTGGTTCATGGTC-3’, SiiD(30-393)-EcoRI-Rev: 5’-GGAGCAGCACT<u>GAATTC</u>TCCGGTAATTACACTGGC-3’, SiiD(43-300)-BamHI-For: 5’-GGAGGAGGAGGA<u>GGATCC</u>GATAATGCTCAGTTAATATCTC-3’, SiiD(43-300)-EcoRI-Rev: 5’-GGAGGAGGAGGA<u>GAATTC</u>TTATGGTTTTATTTCAAAAAGTAAG-3’, SiiD(80-263)-BamHI-For: 5’-GGAGGAGGAGGA<u>GGATCC</u>GATCTGCAAAAAGAATATC-3’, SiiD(80-263)-EcoRI-Rev: 5’-GGAGGAGGAGGA<u>GAATTC</u>TTAATTTATCTGCTTCTCTATT-3’. Purified PCR products were inserted into the pGEX-6P-1 vector (GE Healthcare) using *Bam*HI and *Eco*RI restriction enzymes (New England Biolabs). The correctness of the constructs was verified by DNA sequencing.

A fragment of *siiE* encoding the N-terminal domain and the first two BIg domains was amplified from genomic DNA of *S.* Typhimurium NCTC12023 using primers GST-SiiE-N-For-BamHI: 5’-CGA<u>GGATCC</u>ATACAAAAGTTTTTTGCCGATC-3’ and GST-SiiE-N-Rev-SalI: 5’-AGT<u>GTCGAC</u>TTAGGTGTCGGTTATGATACTATC-3’. The resulting fragment was digested by *Bam*HI and *Sal*I restriction enzymes (New England Biolabs) and ligated to the pGEX-6P-1 vector (pGEX-6P-1-GST::*siiE*_6-429_). SiiE (UniProt accession no. **<u>Q8ZKG6</u>**) variants SiiE(6-236) and SiiE(6-332) were obtained from vector pGEX-6P-1-GST::*siiE*_6-429_ by introducing stop-codons via site-directed mutagenesis using the two-stage PCR protocol [25]. Primers used were SiiE(6-236-stop)-For: 5’-ATAAGCTCGATGCCGAGTCTGTTAAATAATAACTTAAAGTCACATTAGCGCTTGC GGC-3’, SiiE(6-236-stop)-Rev: 5’-GCCGCAAGCGCTAATGTGACTTTAAGTTATTATTTAACAGACTCGGCATCGAGCT TAT-3’, SiiE(6-332-stop)-For: 5’-GTAGCGCCAAACTTGTCATTACTATCGATTCCTAATAAGATAAACCAACATTTGAACTTTCGCCTGAAAG-3’, SiiE(6-332-stop)-Rev: 5’-CTTTCAGGCGAAAGTTCAAATGTTGGTTTATCTTATTAGGAATCGATAGTAATGACAAGTTTGGCGCTAC-3’. Successful mutagenesis was confirmed by DNA sequence analysis.

For the cloning of SiiF (UniProt accession no. **<u>H9L4D0</u>**) variant SiiF(2-120) a fragment of *siiF* was amplified from genomic DNA of *S.* Typhimurium NCTC12023 using primers SiiF-2-For-BamHI: 5’-CTA<u>GGATCC</u>GATAAAAAACTAGAACCTTATTATTTAAG-3’ and SiiF-120-Rev-EcoRI: 5’-CGC<u>GAATTC</u>TCATTATTTTAAATATTCATCTTCAATTTCAAC-3’. The resulting fragment was digested by *Bam*HI and *Eco*RI and ligated to the pGEX-6P-1 vector. SiiF variants SiiF(2-124) and SiiF(2-128) were obtained from vector pWRG549 [5] by PCR using primers SiiF(2-124)/(2-128)-BamHI-For: 5’-CCAGCGGACCA<u>GGATCC</u>GATAAAAAACTAGAACCTTATTATTTAAG-3’, SiiF(2-124)-EcoRI-Rev: 5’-GGAGGACCTGAG<u>GAATTC</u>TTATGCAGATAACTCTTTTAAATATTC-3’, SiiF(2-128)-EcoRI-Rev: 5’-GCACCTCGGGT<u>GAATTC</u>TTATAGTATACTAAATGCAGATAACTC-3’. Purified PCR products were inserted into the pGEX-6P-1 vector using *Bam*HI and *Eco*RI. Correctness of all SiiF constructs was verified by DNA sequencing.

### 2.3. Protein production and purification

GST-tagged SiiD, SiiE and SiiF variants were produced in *E. coli* BL21(DE3) cells (Invitrogen) except for SiiF(2-120), which was produced in BL21(DE3)Star cells (Invitrogen). SiiD(30-393), SiiD(43-300), SiiD(80-263), SiiF(2-124) and SiiF(2-128) were cultured at 37 °C in terrific broth (TB) medium, whereas SiiE(6-236), SiiE(6-332) and SiiF(2-120) were cultured at 37 °C in LB medium. In all cases, 100 μg/mL ampicillin was added to the media. Protein expression of the SiiD variants was induced at an OD_600_ of 0.8 with 0.1 mM isopropyl β-D-1-thiogalactopyranoside (IPTG) and continued at 16 °C for 24 h. SiiE(6-236) and SiiE(6-332) were induced at an OD_600_ of 0.5 with 0.1 mM IPTG, and protein expression continued at 37 °C for 3 h. SiiF(2-120) was induced at an OD_600_ of 0.7 with 1 mM IPTG and protein expression continued at 20 °C overnight, whereas SiiF(2-124) and SiiF(2-128) were induced at an OD_600_ of 0.8 with 1 mM IPTG, and protein expression continued at 20 °C for 24 h. Bacterial cells were harvested by centrifugation, and pellets were disrupted by sonication in PBS buffer (pH 8.0) containing 10 mM EDTA as well as protease inhibitors (Roche) and, in case of the SiiF variants, additionally, 2 mM DTT. The lysate was centrifuged for 1 h at 95,000*g* and the supernatant was loaded onto a Glutathione Sepharose HP (GE Healthcare) column. Fusion proteins were eluted in a buffer containing 50 mM Tris-HCl, pH 8.0, 50 mM NaCl and 10 mM reduced glutathione (the buffer for the SiiF variants contained additionally 2 mM DTT). The GST-tag was cleaved off by adding GST-tagged human rhinovirus 3C (HRV 3C) protease at a mass ratio of 1:200 to the fusion protein and subsequent incubation at 4 °C overnight. HRV 3C protease cleaves the fusion proteins to yield the respective variants with five additional aa residues (Gly, Pro, Leu, Gly and Ser) extending from their N-termini. In case of SiiF(2-128), a size exclusion chromatography run was performed directly after the cleavage to separate the GST-tag and other contaminants from the target. For the other proteins the buffer was exchanged to PBS buffer (pH 8.0) (the buffer for the SiiF variants contained 2 mM DTT in addition) and a second Glutathione Sepharose purification was applied to separate the GST-tag, uncleaved fusion proteins and the HRV 3C protease from the target proteins. In case of SiiD(43-300), a MonoQ 5/50 GL column (GE Healthcare) was required to completely remove the GST-tag. After a buffer exchange to 50 mM Tris-HCl, pH 8.0, 50 mM NaCl, the protein was loaded onto the MonoQ anion-exchange column pre-equilibrated with the same buffer. The protein was eluted with a linear salt gradient ranging from 50 mM to 1 M NaCl. The final purification step for all variants encompassed a size exclusion chromatography run (HiLoad 16/60 or 26/60 Superdex 75 or 200 prepgrade, GE Healthcare) using a buffer with 25 mM Tris-HCl, pH 8.0, 150 mM NaCl (the buffer for the SiiF variants contained 2 mM DTT in addition).

### 2.4. Circular dichroism spectroscopy

To probe the folding of SiiD, SiiE and SiiF variants, circular dichroism (CD) measurements were performed on a J-815 spectropolarimeter equipped with a PTC-423S temperature control element (Jasco). Far-UV spectra were recorded at 20 °C in a 0.1 cm (0.05 cm in case of SiiD(80-263)) quartz cuvette (Hellma) with protein concentrations of 5 µM (SiiD(43-300) and SiiD(80-263)), 7.5 µM (SiiE(6-236)_eth_ and SiiE(6-332)_eth_; eth = variant with ethylated lysines) or 10 µM (SiiF(2-128)) in 10 mM potassium phosphate, pH 7.5 (pH 8.0 for SiiE(6-236)_eth_ and SiiE(6-332)_eth_, 10 mM sodium phosphate, pH 7.5 for SiiF(2-128)), respectively. Spectra were registered with a scanning speed of 20 nm/min. Ten scans per measured sample were averaged and corrected for the sample buffer. The response time was set to 1 s, the bandwidth to 1 nm, the data pitch to 0.1 nm, and the sensitivity to standard. To study the thermal stability of the variants, changes in the ellipticities at 222 nm (SiiD(43-300)), 220 nm (SiiD(80-263)) or 215 nm (SiiF(2-128)) were monitored between 20 °C and 90 °C with a heating rate of 1 °C/min and a response time of 8 s. Normalization of the data to mean residue weight (MRW) ellipticities [θ]MRW was done as described previously [26]. The BeStSel server was used for secondary structure estimations in the wavelength range between 190 nm and 250 nm of the CD spectra [27].

### 2.5. Crystallization

Initial crystallization hits of variant SiiD(43-300) were obtained at 19 °C using the vapor diffusion method with a sitting-drop setup in the MIDAS screen (Molecular Dimensions) condition 1-7 (10% v/v polypropylene glycol 400) with a protein-to-reservoir ratio of 0.2 µL protein (10.5 mg/mL in 25 mM Tris-HCl, pH 8.0) and 0.1 µL of reservoir. The crystals were optimized in a vapor diffusion hanging drop grid screen. In the grid screen the pH of the protein solution (pH 7.7-8.6) versus the polypropylene glycol 400 concentration (5-15%) was evaluated. The best crystals were obtained in a condition with a protein buffer of pH 8.0 and 5% polypropylene glycol 400 in a drop of 2 μL protein and 1 μL reservoir equilibrated against 700 μL of reservoir solution. Crystals appeared after one week and diffracted to 4 Å resolution.

## 3. Results

### 3.1. SiiD

The PAP SiiD links the inner membrane SiiF ABC transporter with the OMP SiiC (Fig. 1A). We performed a sequence search with program HHpred against the Protein Data Bank (PDB) in order to identify structurally homologous PAPs and infer putative domain boundaries in SiiD. Wrongly identified domain boundaries often lead to problems such as reduced protein production yields, enhanced protein aggregation tendency and/or incorrect protein folding. Moreover, the presence of unfolded and disordered segments extending from folded domains also hampers further investigations such as the growth of crystals for X-ray crystallography. The SiiD domain borders were predicted using various bioinformatic tools in combination with the 3D structural information available from the first six HHpred hits, namely LipC from *Serratia marcescens* (PDB entry: 5NEN), EmrA from *Aquifex aeolicus* (PDB entry: 4TKO), Spr0693 from *Streptococcus pneumoniae* (PDB entry: 5XU0), MacA from *E. coli* (PDB entry: 3FPP), AcrA from *E. coli* (PDB entry: 2F1M) and HlyD from *E. coli* (PDB entry: 5C22), all having estimated probabilities of > 99.9% to be structural homologs of SiiD (Supplementary Information S. Fig. 2).

**Figure 1.**
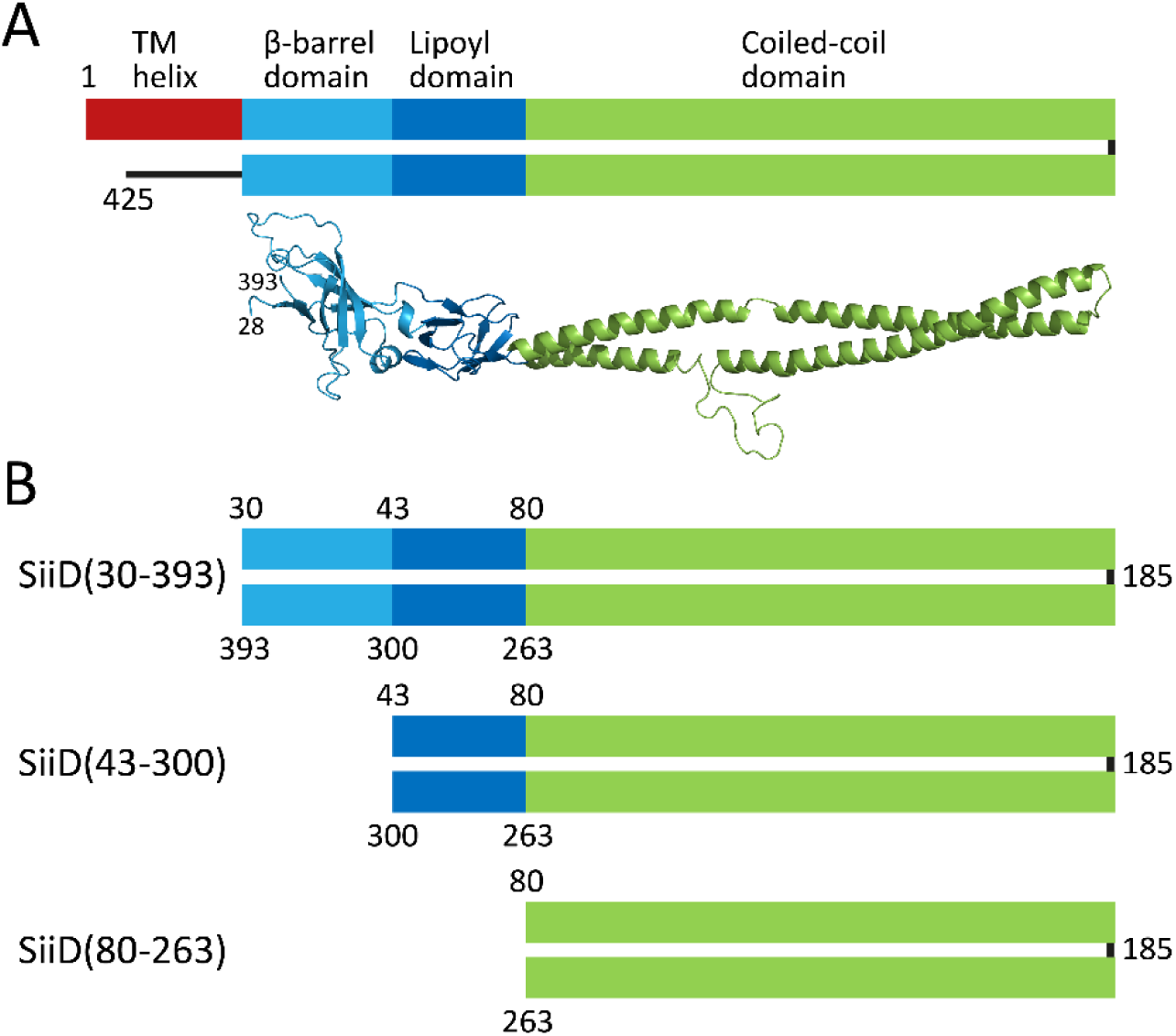
(A) Schematic representation of the SiiD domain structure and SiiD homology model based on the periplasmic adaptor protein EmrA as a template. (B) Designed SiiD expression variants used in this study.

We investigated three different SiiD variants: a first one consisting of the β-barrel, the lipoyl and the coiled-coil domain but without the transmembrane helix, a second one with only the lipoyl and the coiled-coil domain and a third one with the coiled-coil domain, only (Fig. 1B). For the first variant, SiiD(30-393), the domain boundaries of the β-barrel domain had to be determined. Asn30 was chosen as the N-terminal and Gly393 as the C-terminal boundary residue in accordance with a transmembrane helix prediction for SiiD and six structural homologs, a SiiD secondary structure prediction and the known β-barrel domain borders from the structures of EmrA, Spr0693, MacA and AcrA (Supplementary Information S. Fig. 2). The second variant with only the lipoyl and the coiled-coil domain, SiiD(43-300), was selected based on the lipoyl domain boundaries. A SiiD secondary structure prediction in combination with the aligned known lipoyl domains from the six SiiD homologs led to the selection of Asp43 and Pro300 as the N- and C-terminal residues for this variant. The lipoyl domains of all six homologs align very well with each other and with the SiiD lipoyl domain, which is evidenced by a high sequence conservation (Supplementary Information S. Fig. 2). The third variant, SiiD(80-263), was derived from the coiled-coil domain borders. Here, the SiiD secondary structure prediction as well as the aligned known coiled-coil domains from the six SiiD homologs informed the choice of the N- and C-termini for this variant, namely Asp80 and Asn263 (Supplementary Information S. Fig. 2). From the alignment, it can be inferred that the coiled-coil domains of SiiD, LipC, EmrA and HlyD are of similar length, whereas the coiled-coil domains of Spr0693, MacA and AcrA are shorter with roughly half the length.

A homology model of SiiD, based on EmrA and with acceptable quality (QMEAN Z-score = -3.60), is shown in Fig. 1A. The model ranges from Glu28 to Gly393 with the latter residue coinciding with the C-terminal end of variant SiiD(30-393).

The three variants were produced as N-terminal GST-tagged fusion proteins. While SiiD(30-393) and SiiD(80-263) were purified in three chromatographic steps, a GST affinity column, a second GST affinity column after HRV 3C protease treatment and a final size exclusion column (Fig. 2A, C, E), SiiD(43-300) required an additional anion exchange chromatography step after the second GST affinity column to remove any remaining GST protein. Highly pure protein in mg amounts was obtained for all three SiiD variants according to SDS gel analyses (Fig. 2B, D, F). Interestingly, the yields varied considerably between the different variants. SiiD(43-300) exhibited the best overproduction, followed by SiiD(80-263) and finally SiiD(30-393).

**Figure 2.**
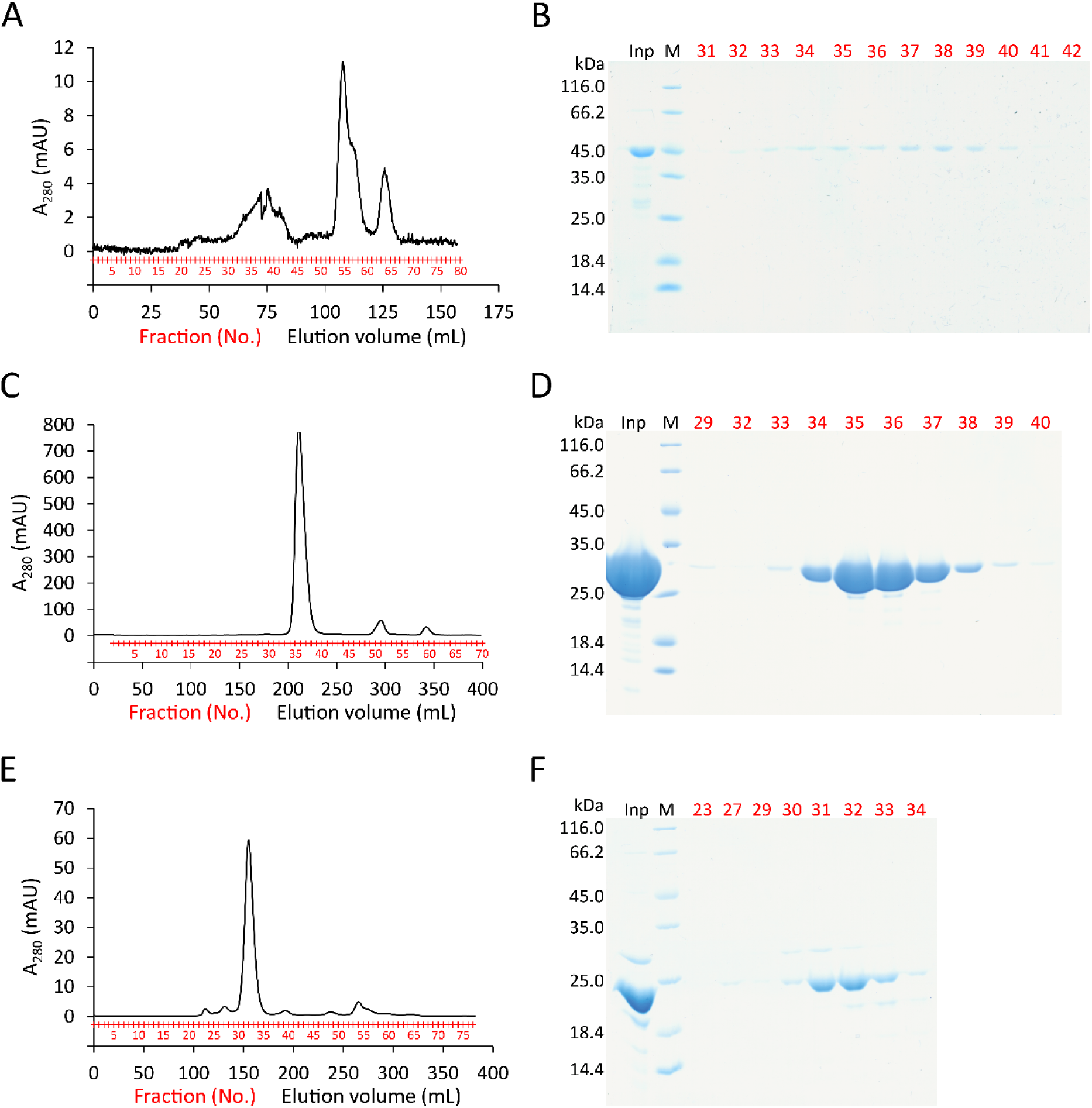
(A) Elution profile (HiLoad 16/60 Superdex 200 size exclusion chromatography) and (B) SDS-PAGE of the final purification step of SiiD(30-393). (C) Elution profile (HiLoad 26/60 Superdex 200) and (D) SDS-PAGE of the final purification step of SiiD(43-300). (E) Elution profile (HiLoad 26/60 Superdex 75) and (F) SDS-PAGE of the final purification step of SiiD(80-263). The SDS-PAGE lanes are labeled with Inp for input sample, M for protein size standard with molecular weights on the left. Numbers in red indicate the start of each eluate fraction.

Proper folding of SiiD(43-300) and SiiD(80-263) was experimentally investigated by CD measurements (Fig. 3A, B). The spectra of both proteins are typical for proteins containing α-helical structure with minima at 209 nm and 222 nm as well as a maximum at 192 nm. Estimation of the secondary structure content with the BeStSel server (Table 1) yielded values that are close to those derived from the SiiD homology model, except for the β-sheet content, which is 0% in SiiD(43-300) (BeStSel) and 13.2% in the model. We can only speculate about the reason for this discrepancy - potentially the lipoyl domain adopts a β-sheet architecture, which is currently not recognized by the BeStSel server. To investigate the thermal stability, both variants were subjected to a heat denaturation. The resulting melting temperatures are 47.8 °C ± 0.1 °C for SiiD(43-300) and 51.7 °C ± 0.1 °C for SiiD(80-263). The obtained spectra suggest cooperative unfolding of the proteins with possibly aggregation behavior above 60 °C (Supplementary Information S. Fig. 3).

**Figure 3.**
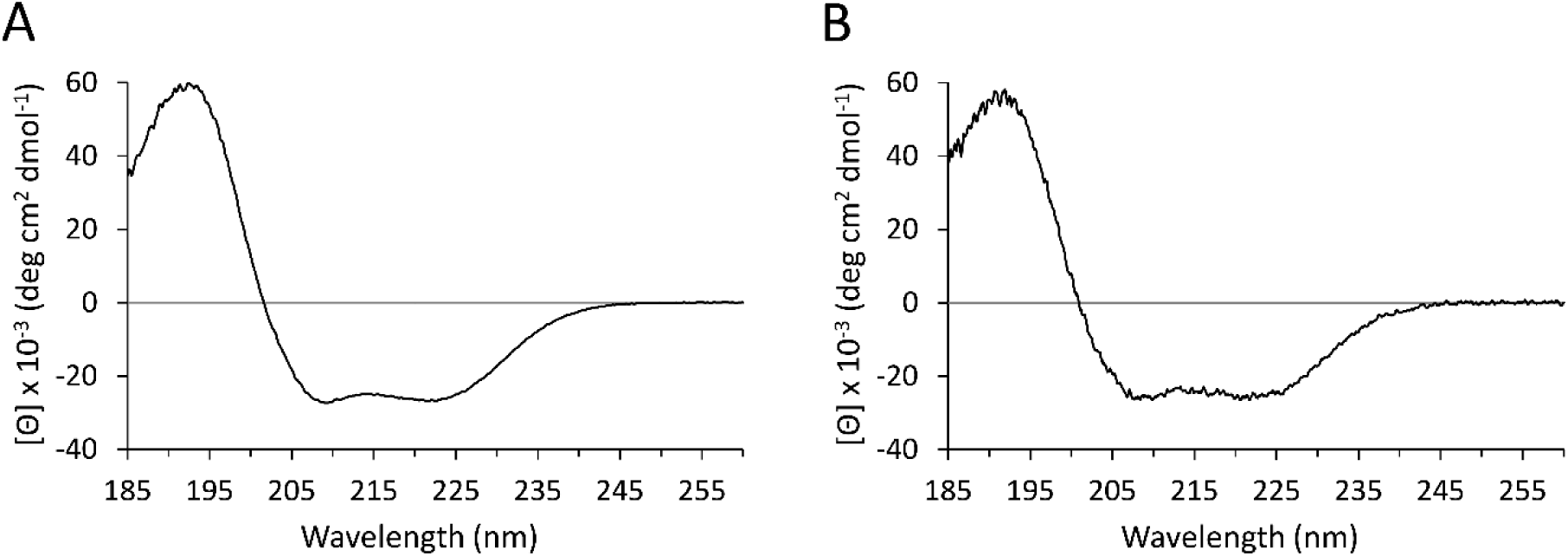
Circular dichroism (CD) spectra of (A) SiiD(43-300) and (B) SiiD(80-263) in the far UV region.

**Table 1.** Secondary structure content estimations.

|  | SiiD(43-300) | SiiD(80-263) | SiiE(6-236) | SiiE(6-332) | SiiF(2-120) | SiiF(2-128) |
| --- | --- | --- | --- | --- | --- | --- |
| **BeStSel server** |  |  |  |  |  |  |
| Helix (%) | 68.7 | 65.2 | 24.1 | 15.6 |  | 20.1 |
| β-sheets (%) | 0.0 | 0.0 | 17.4 | 24.9 | | 30.2 |
| Turns (%) | 8.7 | 9.4 | 14.1 | 14.3 |  | 11.9 |
| Others (%) | 22.6 | 25.4 | 44.4 | 45.2 |  | 37.8 |
| **Homology model<sup>a</sup>** |  |  |  |  |  |  |
| Helix (%) | 55.0 | 76.6 |  |  | 45.0 | 26.0 |
| β-sheets (%) | 13.2 | 0.0 | | | 12.5 | 24.4 |
| Turns (%) | 10.5 | 6.5 |  |  | 11.7 | 7.9 |
| Others (%) | 21.3 | 16.9 |  |  | 30.8 | 41.7 |
| **JPred4 server** |  |  |  |  |  |  |
| Helix (%) |  |  | 30.9 | 22.0 |  |  |
| β-sheets (%) | | | 24.6 | 31.0 | | |
| Turns and Others (%) |  |  | 44.5 | 47.0 |  |  |

<sup>a</sup> Values were calculated from the corresponding homology models using the DSSP program [22]. The assignment of DSSP secondary structure elements followed the scheme utilized in [27] (H: helix, E: $\beta$ -sheets, T: turns and G/S/I/B/without letter: others). The SiiF(2-120) homology model was obtained from program MODELLER [33] implemented in the HHpred server [17] with the structure of RsmG from *Bacillus subtilis* (PDB entry: 1XDZ) as a template.

Protein crystals were obtained with SiiD(43-300) under various conditions, and the best crystals are shown in Fig. 4A. These crystals did not diffract to beyond 4 Å resolution (Fig. 4B), and any attempts to increase the diffraction quality, using an additive screen or crystal dehydration or protein that was chemically modified by lysine methylation, did not improve diffraction properties.

**Figure 4.**
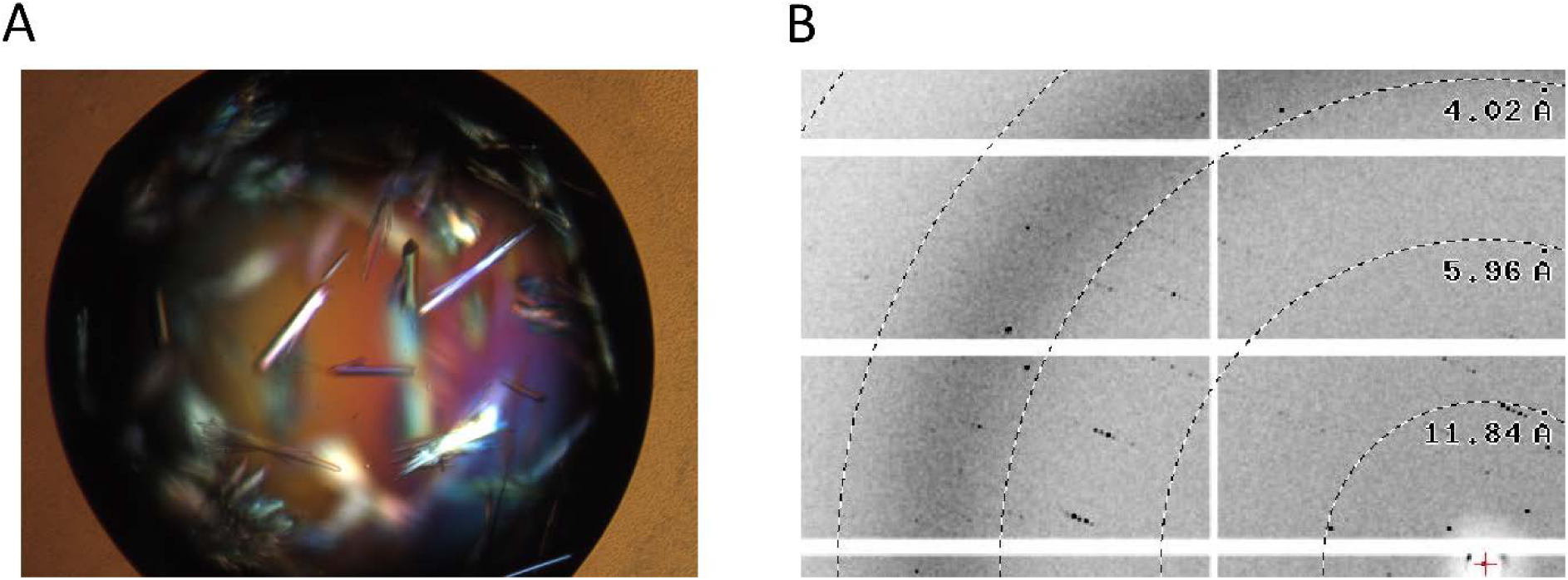
(A) Crystals of SiiD(43-300) and (B) diffraction pattern of a SiiD(43-300) crystal with spots to approximately 4 Å resolution.

### 3.2. SiiE N-terminal region

The 5559-residue-long giant adhesin SiiE (Fig. 5A) is the substrate of the SPI4-T1SS. Its long filamentous structure (about 175 nm [16]) makes it very unlikely that crystals, suitable for X-ray structure determination, can be grown of the whole entity. Crystallization trials with an engineered deletion variant where the N-terminal half of BIg2 was fused to the C-terminal half of BIg49 resulting in a variant containing aa 5-403 and aa 5053-5559 (mini-SiiE) were unsuccessful until now. Limited proteolysis of mini-SiiE identified a stable fragment, SiiE-BIg50-52, which readily crystallized [28] and finally led to its high resolution 3D structure [14]. With the aim to gain structural insight into the N-terminal region, two variants were selected for protein production. The choice of the respective N- and C-termini for these variants relied entirely on bioinformatic analyses from Gerlach *et al.* [12] and Wagner *et al.* [16] since a search with HHpred did not find structural homologs with high likeliness in the region aa 1-240. SiiE’s N-terminal domain is made up of three parts, aa 1-116 with predicted β-sheet secondary structure, aa 117-172 with predicted coiled-coil structure containing eight heptad repeats and aa 173-236 with again predicted β-sheet structure, although disorder predictions point to a possible complete disorder of the region encompassing aa 173-236. Two variants were designed, SiiE(6-236) comprising the N-terminal domain, as well as SiiE(6-332) comprising the N-terminal domain and the first BIg domain (Fig. 5B).

**Figure 5.**
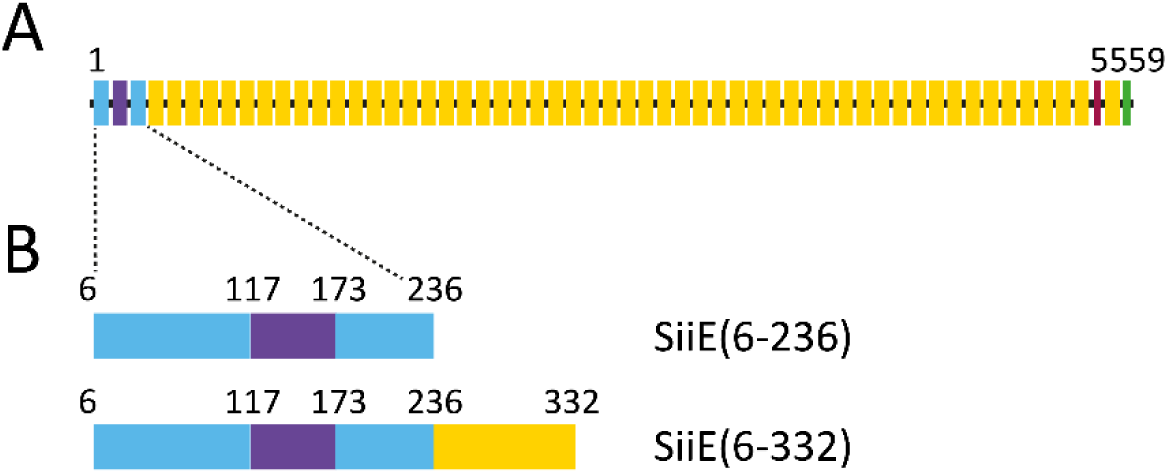
Schematic representations of (A) full length SiiE and (B) of the expression variants used in this study. The N-terminal domain (aa 1-236) comprises two predicted β-sheet regions (light blue) with a coiled-coil region (purple) in between. The BIg-domains are colored yellow. At the C-terminus an insertion (red) between BIg52 and BIg53 and a secretion signal (green) is located.

Both variants were produced as N-terminal GST-tagged fusion proteins and purified in three chromatographic steps including tag removal. Highly pure protein was obtained according to SDS gel analyses (Fig. 6) with considerably larger yields for SiiE(6-236) compared to SiiE(6-332). For crystallization trials, the proteins were chemically modified by lysine ethylation (data not shown). CD measurements were performed using these ethylated (eth) variants (SiiE(6-236)_eth_ and SiiE(6-332)_eth_). The analysis showed that both variants are properly folded with spectra typical for proteins containing both α-helical and β-sheet structure in accordance with the predicted coiled-coil and β-sheet structure (Fig. 7) and the estimation of the secondary structure content (Table 1). The spectrum from SiiE(6-236)_eth_ exhibits a maximum at 190 nm and two minima at 206 nm and 222 nm, the spectrum from SiiE(6-332)_eth_ a maximum at 191 nm and two minima at 207 nm and 221 nm. The spectrum from SiiE(6-332)_eth_ shows a zero crossing shifted to a longer wavelength compared to the SiiE(6-236)_eth_ spectrum, which can be attributed to a higher beta sheet fraction caused by the additional BIg domain in SiiE(6-332). Heating the samples up to 96 °C does not allow for the monitoring of the unfolding of the protein variants but is only accompanied by slight decreases in ellipticities (recorded at 220 nm, data not shown). At 75 °C the CD signal for SiiE(6-332)_eth_ decreases slightly more distinctly, which might indicate aggregation of the protein molecules.

**Figure 6.**
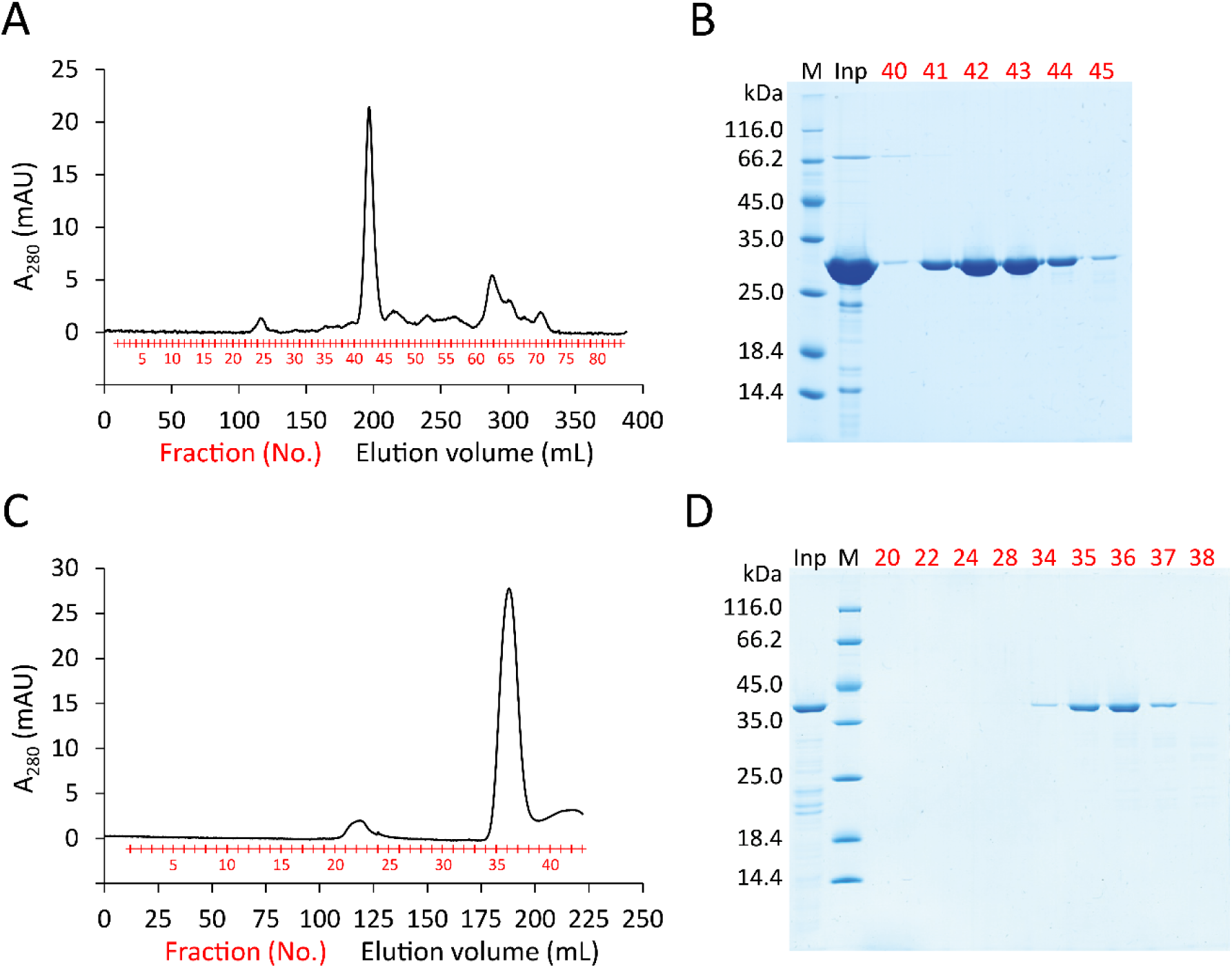
(A) Elution profile (HiLoad 26/60 Superdex 75 size exclusion chromatography) and (B) SDS-PAGE of the final purification step of SiiE(6-236). (C) Elution profile (HiLoad 16/60 Superdex 200) and (D) SDS-PAGE of the final purification step of SiiE(6-332). The SDS-PAGE lanes are labeled with Inp for input sample, M for protein size standard with molecular weights on the left. Numbers in red indicate the start of each eluate fraction.

**Figure 7.**
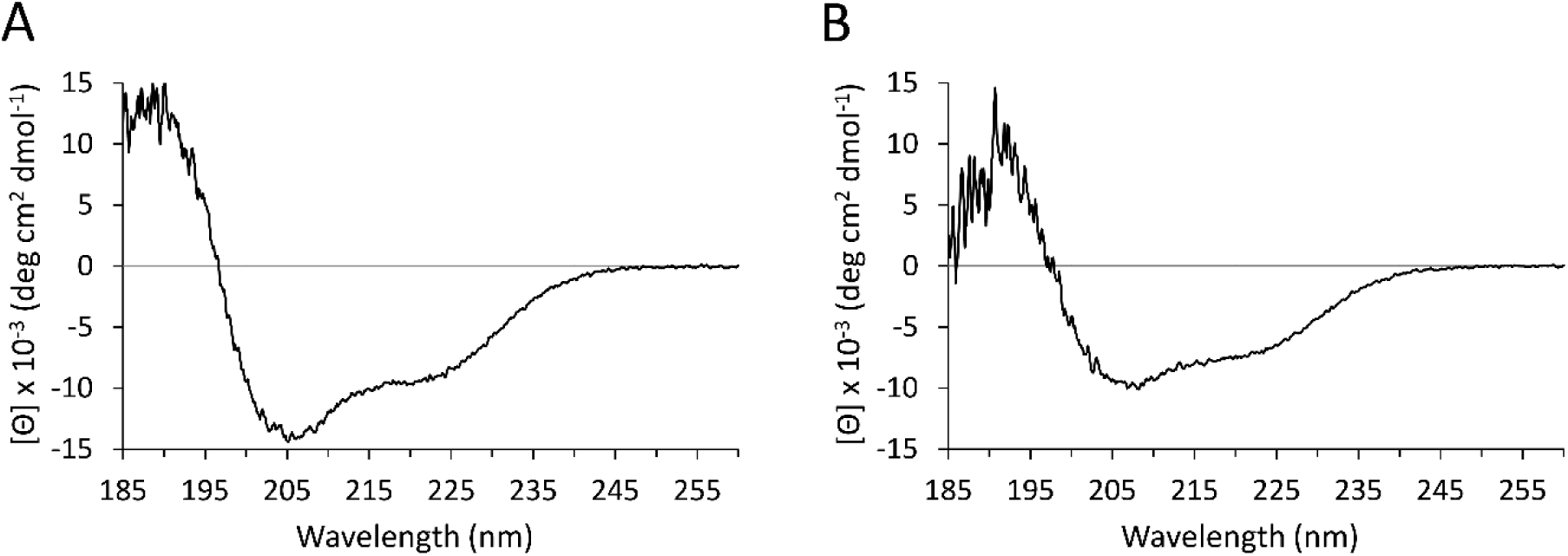
Circular dichroism (CD) spectra of (A) SiiE(6-236)_eth_ and (B) SiiE(6-332)_eth_ (eth = lysines are ethylated) in the far UV region.

### 3.3. SiiF N-terminal domain

The anticipated domain architecture of the ABC transporter SiiF is depicted in Fig. 8A. To gain insight into the possible 3D structure of the SiiF N-terminal domain, we initially performed a search with the program HHpred against the PDB using the N-terminal 150 aa residues, only. The proteins with the highest probability (> 98%) for being structurally homologous to SiiF are HlyB from *E. coli* (PDB entry: 3ZUA), ComA from *Streptococcus mutans* (PDB entry: 3K8U), a toxin secretion ATP-binding protein from *Vibrio parahaemolyticus* (PDB entry: 3B79) and PCAT1 from *Clostridium thermocellum* (PDB entry: 4RY2). Sequence identities to the SiiF N-terminal domain are low, namely 14%, 11%, 11% and 11%, respectively. All four proteins belong to the family of ABC transporters with ComA and PCAT1 exhibiting an N-terminal C39 peptidase domain, whereas HlyB and the toxin secretion ATP-binding protein exhibit an N-terminal CLD without the catalytically essential cysteine. The overall structure of the C39 peptidase domain and the CLD is similar comprising a six-stranded central β-sheet surrounded by five α-helices. However, a subsequent database search using PSI-BLAST [29] revealed a higher sequence identity between the SiiF N-terminal domain and the N-terminal half of RsmG, a ribosomal RNA methyltransferase (212 aa residues) from *Muricauda ruestringensis*, namely 26%. The latter differs considerably in structure and function from the CLDs and C39 peptidase domains.

**Figure 8.**
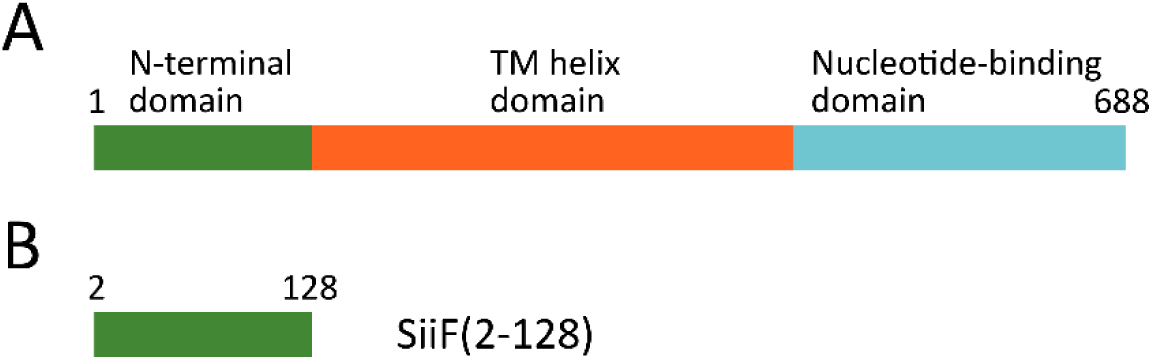
Schematic representations of the (A) SiiF domain structure and (B) one of the designed expression variants - SiiF(2-128).

Based on these bioinformatic surveys, three variants were chosen for the production of the SiiF N-terminal domain: SiiF(2-120), SiiF(2-124) and SiiF(2-128) (Fig. 8B). The SiiF(2-120) variant originates from the predicted homology to the N-terminal half of RsmG from *M. ruestringensis*. Because there is currently no 3D-structure available of the *M. ruestringensis* protein, the structure of RsmG from *Bacillus subtilis* (PDB entry: 1XDZ) served as a template for modeling the *M. ruestringensis* structure first (data not shown). According to program HHpred, RsmG from *B. subtilis* has an estimated probability to be structurally homologous to RsmG from *M. ruestringensis* of 99.56% and the sequence identity adds up to 32%. Using the model of RsmG from *M. ruestringensis* in combination with the alignment between RsmG from *M. ruestringensis* and the SiiF N-terminal domain (data not shown), Lys120 in SiiF was identified as a residue localized at the beginning of the C-terminal half of RsmG and hence chosen as the terminal residue for the variant SiiF(2-120). The SiiF(2-128) variant was derived from the predicted homology to CLDs and C39 peptidase domains. Program SWISS-MODEL generated a homology model of full-length SiiF with PCAT1 (PDB entry: 4RY2) as the best template. From the underlying sequence alignment (Supplementary Information S. Fig. 4) it can be deduced that SiiF’s N-terminal domain would belong to the CLDs as the catalytically essential cysteine of C39 peptidases (Cys21 in PCAT1 [30]) is absent in SiiF. In PCAT1 the last secondary structure element of the N-terminal C39 peptidase domain is a β-strand ranging from Gly135 to Leu138. Since Leu128 in SiiF is predicted to be the last residue of the analogous β-strand, we have chosen Leu128 as the terminal residue for the variant SiiF(2-128). Finally, a variant was designed, that accounts for the uncertainty of the exact domain boundary of the SiiF N-terminal domain. For this purpose a variant ending in between aa residues 120 and 128 - SiiF(2-124) - was selected.

All three variants were produced as N-terminal GST-tagged fusion proteins and purified in three subsequent chromatographic steps including tag removal. While SiiF(2-120) and SiiF(2-124) yielded almost exclusively aggregated protein in the final size exclusion chromatography step, highly pure protein in mg amounts was obtained for SiiF(2-128) as judged from a SDS gel analysis (Fig. 9A, B). To determine the oligomerization state of SiiF(2-128), a molecular weight analysis using a calibrated Superdex 75 Increase 10/300 GL column (GE Healthcare) was performed. The protein’s experimentally determined molecular weight was 21.5 kDa, which is 1.46 times higher than the theoretical molecular weight of 14.7 kDa, suggesting a monomeric state (Supplementary Information S. Fig. 5).

**Figure 9.**
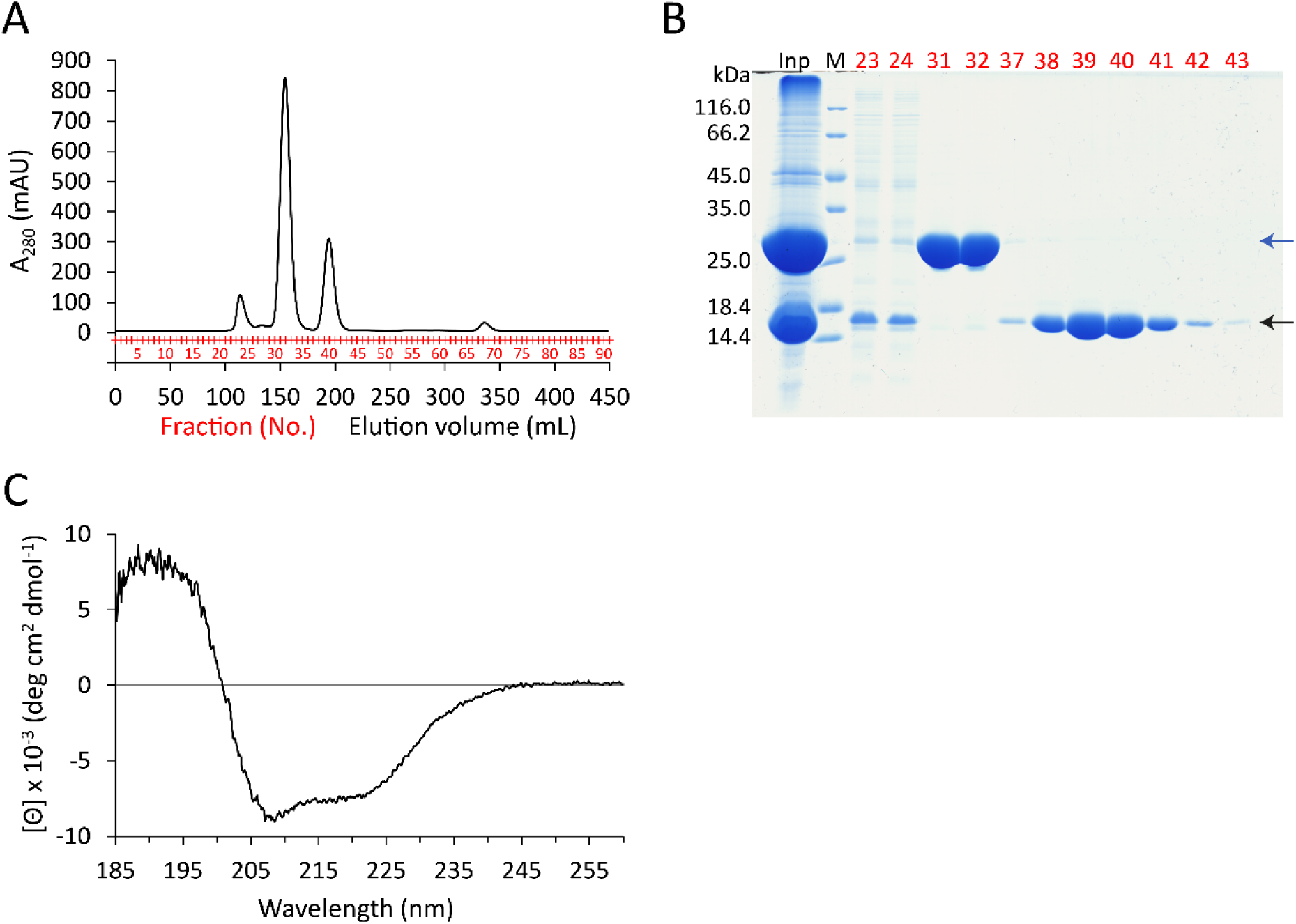
(A) Elution profile (HiLoad 26/60 Superdex 75 size exclusion chromatography) and (B) SDS-PAGE of the final purification step of SiiF(2-128). The SDS-PAGE lanes are labeled with Inp for input sample, M for protein size standard with molecular weights on the left. Numbers in red indicate the start of each eluate fraction. Blue arrow: GST-tag. Black arrow: SiiF(2-128). (C) Circular dichroism (CD) spectrum of SiiF(2-128) in the far UV region.

CD analysis showed that SiiF(2-128) is folded and contained both α-helical and β-sheet structure (Fig. 9C). The CD signal is positive below 200 nm with a maximum at approximately 190 nm and negative above 200 nm with two minima at around 208 nm and 222 nm. The values of the estimated secondary structure content from the BeStSel server are in clearly better agreement with those derived from the SiiF homology model that was based on the PCAT1 structure, compared to those derived from the RsmG structure as the model template (Table 1). In a thermal scanning CD measurement SiiF(2-128) exhibited a decrease in ellipticity upon heating, suggesting that instead of a thermally induced unfolding aggregation occurs (Supplementary Information S. Fig. 6).

## 4. Discussion

*Salmonella* infections constitute severe health threats for humans. The SPI4-encoded T1SS formed by the proteins SiiA to SiiF initiates the first contact between the *Salmonella* bacterium and the host cell. Currently, neither a cryo-electron microscopy nor an electron tomography structure is available of this T1SS. At the same time, only one single crystal structure has been determined so far. The structure describes a portion of one of the six proteins that are encoded by the SPI4, namely BIg domains 50 to 52 of the 5559-residue-long secreted adhesin SiiE [14]. Some information with regard to the mechanism of the SPI4-encoded T1SS can be inferred from knowledge available on homologous secretion systems such as, for example, the *E. coli* HlyA T1SS [11]. However, current progress in understanding the particularities of the *S.* Thyphimurium SPI4-encoded T1SS is hampered by a scarcity of experimental data on structure-function relationships in the SPI4-encoded T1SS and its protein constituents.

In an initial effort to overcome the data scarcity, we focused on the recombinant protein production of domains from three of the six proteins of the T1SS. These proteins were chosen for the following reasons: SiiD constitutes the PAP of the T1SS. Its function is to bridge between the SiiF ABC transporter located within the inner membrane and the OMP SiiC located in the outer membrane. By virtue of bridging the space between the inner and outer membrane, SiiD spans across the peptidoglycan layer. Interestingly, of the two additionally SPI4-encoded proteins SiiA and SiiB, the protein SiiA contains a periplasmic peptidoglycan-binding domain (data not shown). Thus, we hypothesized that the colocalisation of SiiD and the peripasmic domain of SiiA could provide for a functional link between the canonical T1SS proteins SiiF, SiiD and SiiC and the auxillary proteins SiiA and SiiB. This link could originate from either a direct or peptidoglycan-mediated interaction between SiiD and SiiA, and both of these possibilities could be explored by a direct interaction assay.

Similar hypotheses also led to the selection of the N-terminal domains of SiiF and SiiE for recombinant protein production. Whereas the N-terminal domain of SiiF shares low sequence homology to domains with known three-dimensional structure, no homology could be detected at all in case of the N-terminal domain of SiiE. Moreover, we hypothesized that the N-terminal domain of SiiF might play an important role in initiating the threading of the secreted adhesin SiiE across the T1SS, possibly, by specifically interacting with SiiE, a behavior seen in other T1SSs before [8]. Likewise, it is also possible that this domain specifically interacts with the N-terminus of SiiE and is involved in the retention of SiiE on the bacterial surface. To test whether a direct interaction between the N-terminal domains of SiiF and SiiE occurs, both proteins were recombinantly produced, first.

In almost all cases, the choices of different domain boundaries led to drastic differences in protein production yields. Since *Salmonellae* are genetically close to *E. coli*, we propose that differences in production yields reflect differences in the stability of the translated protein products and are not caused by indirect effects, such as, for example, differences in codon usage. Codon usage is highly similar in *E. coli* and *S.* Typhimurium and, therefore, should not alter protein translation levels in variants with different gene sequences [31].

In all three proteins studied here, at least one construct could be identified that yielded high amounts of protein upon heterologous expression. These successes provide a direct experimental validation for the domain boundaries of these protein variants that were initially predicted from bioinformatics considerations, only. It is interesting that, for example, in case of SiiF, construct differences at the C-terminus as small as four residues led to drastic differences in recombinant protein overproduction rates. Whereas variants SiiF(2-120) and SiiF(2-124) yielded almost exclusively aggregated protein samples, variant SiiF(2-128) can be overproduced with high yields and behaves as a monomer when investigated on a size exclusion chromatography. Together with the estimation of the secondary structure contend inferred from CD investigations, these results clearly favor the SiiF model that was generated based on the predicted homology to CLDs and C39 peptidase domains over a model that was generated based on the observed higher sequence identity to the N-terminal half of methyltransferase RsmG. Thus, the N-terminal domain of SiiF forms a separate unit that might fulfill a function that can be clearly distinguished from the sole translocation of substrates across membranes as performed by ABC transporters consisting of only the canonical nucleotide binding and transmembrane domains. In view of the absence of the catalytically essential cysteine of C39 peptidases, the function of SiiF’s N-terminal domain might be related to the function exhibited by CLDs, which are essential in recruiting the substrate to the transporter [8].

In case of SiiE, both variants, SiiE(6-236) comprising the N-terminal domain and SiiE(6-332) comprising the N-terminal domain and the first BIg domain, yielded properly folded protein according to CD analyses. These results indicate that the domain boundaries of these variants were chosen adequately and that the N-terminal domain forms a unit that can be expressed separately. Secondary structure and coiled-coil structure predictions indicate that the N-terminal domain consists of a central coiled-coil region flanked by two β-sheet regions. For the first BIg domain solely β-sheet structure is predicted. The estimation of the secondary structure content for SiiE(6-236) and SiiE(6-332) deduced from their respective CD spectra is in agreement with these predictions. Coiled-coils mediate the homo- or heteromeric assembly of protein subunits and the coiled-coil region in SiiE’s N-terminal domain was postulated to play an essential role in the retention of SiiE [16].

In case of SiiD, three variants were designed in order to increase the chances for obtaining soluble protein with high yields. Most of the published PAP 3D structures comprise all three periplasmic domains, the β-barrel, the lipoyl and the coiled-coil domain. In contrast, for the HlyD-structure determination, a variant with an acceptable production level was identified that covers only the lipoyl and the coiled-coil domain [32]. Interestingly, we observe considerable variations in production levels between the different SiiD variants with the best overproduction for SiiD(43-300), which also comprises only the lipoyl and the coiled-coil domain. On the other hand, the yield was much lower for variant SiiD(30-393) that contains all three domains possibly implicating that the chosen N- and C-terminal borders of the β-barrel domain were not ideal. CD spectroscopy measurements verify the correct folding of SiiD(43-300) and SiiD(80-263), comprising only the coiled-coil domain, and both variants unfold cooperatively at around 50 °C when investigated in a CD thermal stability analysis.

First experiments aiming at probing the interaction between the produced protein domains remained so far inconclusive. This was also the case when probing interactions with additional partners such as the peptidoglycan-binding domain of SiiA or the C-terminal portion of SiiE (data not shown). Structural studies have been initiated, and in case of SiiD initial crystals have been obtained. No crystals have so far been obtained with SiiF(2-128), SiiE(6-236) and SiiE(6-332). The size of the SiiF(2-128) protein renders SiiF also amendable to a structure determination *via* NMR. Detailed structural and functional studies will be a prerequisite for better understanding the particularities of this remarkable T1SS. In the long run, these studies might upon up new routes for combatting *Salmonella* infections.

## Acknowledgments

We acknowledge the Helmholtz-Zentrum Berlin for the allocation of synchrotron radiation beamtime at beamlines 14.1 and 14.2 at BESSY II; and thank Dr. Manfred Weiss for help with data collection at BESSY II. We also want to thank Katharina Hohl and Kerstin Hof for technical assistance.

