## Supplementary material for "Recombinant protein production and purification of SiiD, SiiE and SiiF - components of the SPI4-encoded type I secretion system from *Salmonella* Typhimurium"

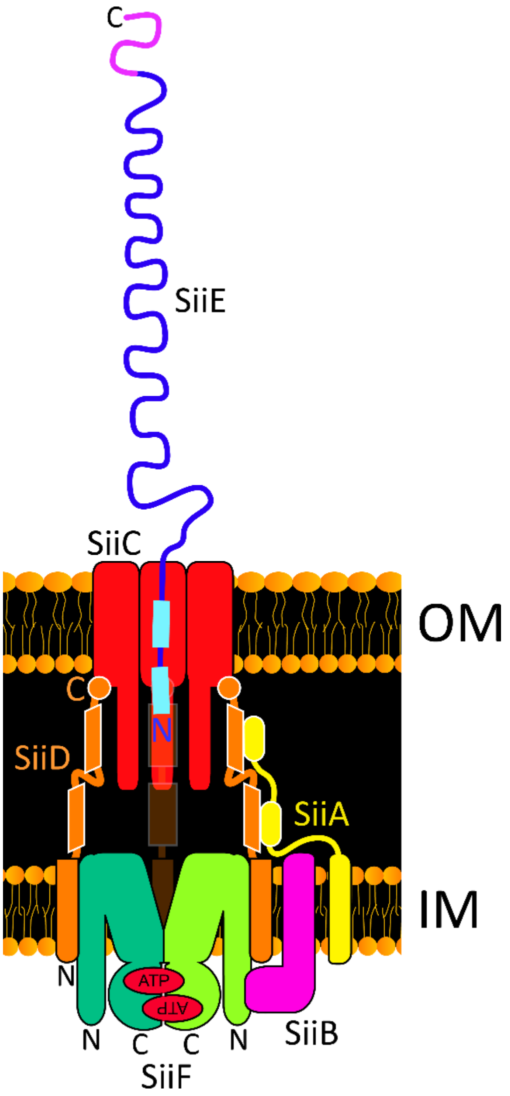


**S. Fig. 1.** Architecture of the *Salmonella* Pathogenicity Island 4 encoded type I secretion system. SiiA and SiiB (potential proton channel proteins), SiiC (outer membrane protein channel), SiiD (periplasmic adapter protein), SiiE (adhesin), SiiF (ABC transporter). N- and C-termini of SiiD, SiiE and SiiF are labelled. OM: outer membrane, IM: inner membrane.

```
(SiiD)                                                              1                 10         20                           30
(SiiD)                                                              ----HHHHH          HHHHHHH HHHHHHHHH                   H----EEEEEEEE
SiiD      ----------------------------------------------------MNRRQSDHL----------MMIIISL-TILIIILTY-------------------FIEINSVVHGQGV
LipC      ------------------------MSTHIGEPQDS-----------YTEEIPQDERRFTRM----------GWLVVGIGLFGFLAWAA-------------------FAPLDKGVASPGS
EmrA      ---------------------------------------------------MKKYIGVFTV----------VALIVGFGIYAFFFIKH-------------------RIEY---AITNAV
Spr0693   ---------------------------------------------------MKKKNGKAKKWQLYAAIGAASVVILGAGGILLFRQPSQTA----LKDEPIHLVVAKEGSVASSVLLSGT
MacA      ---------------------------------------------------MKKRKTVKKRYVI-------ALVIVIAGLITLWRILN-------APVPTYQTLIVRPGDLQQSVLATGK
AcrA      ---------------------------------------------------MNKNRGF---TPL-------AVVLMLSGSLALTGCDDKQAQQGGQQMPAVGVVTVKTEPLQITTELPGR
HlyD      MKTWLMGFSEFLLRYKLVWSETWKIRKQLDTPVREKDENEFLPAHLELIETPV-SRRPRLV----------AYFIMGFLVIAFI-LSV-------------------LGQVEIVATANGK
Cons.     ---------------------------------------------------0413441000----------55798541576860511-------------------13460017155+3

(SiiD)    40         50        60        70        80        90         100       110           120          130           140
(SiiD)    EEE---EEEEE -----EEEEEE-----EEE---EEEEE----HHHHHHHHHHHHHH  HHHHHHHHHH----------    ---HHH   HHHHHHHHHHH HHHHHHHH   H
SiiD      ITTKDNAQLIS-LSKGGTIQDIYVAEGDTVKKGELLAKVVNLDLQKEYQRYRTQKGY--LDKDVNEISFILDKENESGL----ITLDGT---RSLSNKEVKAN-IELVHSQI---R----
LipC      VTVSGNRKTVQ-APASGIIKNIAVRDGDKVKAGEVLVQLSQVQAQAQV--DSLRDQY--YTTLATEGRLLAERDGLSIV-------TFSPILDAVKDKPRVAEIIALQTQLFAS-RRQAL
EmrA      FVKADELSYLS-FRVSGKVIEVYKDLGDYVKRGEALAKLDPTYYELEKRTLEKKMSALLEKKKALEIKIQKLEKGLHIS---------------LSAK----------------------
Spr0693   VTAK-NEQYVYFDASKGDLDEILVSVGDKVSEGQALVKYSSSEAQAAYD--SASRAVARAD--------------RHIN-----------------------------------------
MacA      LDAL-RKVDVG-AQVSGQLKTLSVAIGDKVKKDQLLGVIDPEQAENQIK--EVEATLMELRAQ-----RQQAEAELKLA-----------------------------------------
AcrA      TSAY-RIAEVR-PQVSGIILKRNFKEGSDIEAGVSLYQIDPATYQATYD--SAKG-------D-----LAKAQAAANIA-----------------------------------------
HlyD      LTLSGRSKEIK-PIENSIVKEIIVKEGESVRKGDVLLKLTALGAEADT--LKTQSSL--LQARLEQIRYQILSRSIELNKLPELKLPDEPYFQNVSEEE-VLRLTSLIKEQFSTWQNQKY
Cons.     6442-623393-12248393544434*63943545*546453358344---444301--01-0-----00001002375-----------------------------------------

(SiiD)                  150       160       170       180       190       200       210              220       230       240
(SiiD)    H       HHHHHHHHHHHHHHHHHHHHHHHHHHHHHHHHHHHHHHHHHHHHHHHHHHHHHHHHHHHHHHHHHHHHHHHHH       HHHHHHHHHHHHHHHHHHHHHHHHHHHHHHHHH
SiiD      A-------KELKKTSLDSEISGLQEKLSSKEKELALLAEEINILSPLVKKGISPYTNFLNKKQAYIKVKSEINDIESSIT-------LKKDDIELVVNDIEALNNELRLSLSKIISKNLQ
LipC      QSEIDGYKQSMDGIR--FQLKGLQDSRGNKQIQLSSLREQMNSMKQLAADGYLPRNRYLEVQRQFAEVNSSIDETVGRIGQLQKQLLESQQRIDQRFADY---QREVRTQ----LAQTQM
EmrA      ---------KLKVESLKKKREALREKLLQVEEKIKLVKLDWERYKSLFQKGLIPRRKFEEVDTNLKVLLHEREYLEKSIQEINTEIKRAKKGIENARNEFKTIE-----ELKKELSSLEE
Spr0693   ---------------------ELNQARNEAASA-----PAPQLPAPVGGEDAT-------VQSPTPVAGNSVASIDAQLGDARDARADAA------------------------------
MacA      ---------------------RVTYSRQQR----------------LAQTKAV-------SQQDLDTAATEMAVKQAQIGTIDAQIKRNQ------------------------------
AcrA      ---------------------QLTVNRYQK----------------LLGTQYI-------SKQEYDQALADA-------QQANAAVTAAK------------------------------
HlyD      QKELNLDKKRAERLTILARINRYENVSRVEKS-------RLDDFRSLLHKQAIAKHAVLEQENKYVEAANELRVYKSQLEQIESEILSAKEEYQLVTQLF---KNEILDK----LRQTTD
Cons.     ---------------------365333232----------------9635355-------355261373363001001241110001244------------------------------

(SiiD)     250       260       270       280       290         300       310       320       330       340       350
(SiiD)    HHHHHHHHHHHHHHH--HEEEEE----EEEEEEEEE----EEE-----  EEEEE----EEEEEEEE------EEE----EEEEEEE-------EEEEEEEEE----         ---
SiiD      ELEVVNSTLKVIEKQINEEDIYSPVDGVIYKINKSATTHGGVIQAADL--LFEIKPKVRTMLADVKILPKYRDQIYVDEAVKLDVQSIIQPKIKSYNATIDNISPDS---------YEE-
LipC      DASEFRNKLQMADFDLGNTAITSPVDGTVVGLN--IFTQGGVVGAGDH--LMDVVPSQATLVVDSRLKVDLFDKVYNGLPVDLMFTAFNQNKTPKIPGTVTLVSADR---------LVD-
EmrA      EIKSLKERIKTAEQKIKDTVLIAPFDGVVAKR---FISRGDVVRAGQP--AFALVNPES-FYVEVLLEETKLKGVKVGNKAYVRLDAYPDILF---EGVVEEISPAS---------AATF
Spr0693   ------AQLSKAQSQLDATTVLSTLEGTVVEVN---SNVSKSPTGASQ--VMVHIVSNENLQVKGELSEYNLANLSVGQEVSFTSKVYPDKKW---TGKLSYISDYP------KNNGEAA
MacA      ------ASLDTAKTNLDYTRIVAPMAGEVTQIT---TLQGQTVIAAQQAPNILTLADMSAMLVKAQVSEADVIHLKPGQKAWFTVLGDPLTRY---EGQIKDVLPTP-------------
AcrA      ------AAVETARINLAYTKVTSPISGRIGKSN---VTEGALVQNG-QATALATVQQLDPIYVDVTQSSNDFLRLKQELA------NGTLKQE---NGKAKVSLITSDGIKFPQDGTLEF
HlyD      SIELLTLELEKNEERQQASVIRAPVSGKVQQLK--VHTEGGVVTTAET--LMVIVPEDDTLEVTALVQNKDIGFINVGQNAIIKVEAFPYTRYGYLVGKVKNINLDA---------IED-
Cons.     ------33945551573352938774*394331---252835637+12--5834542431838563622223239454422010006253353---3+474355233---------100-

(SiiD)             360       370           380       390       400           410                   420
(SiiD)          ------EEEEEEEEEE---    -----------EEEEEEEE---HHHHHHHHHHHH    HHHH            H-------------
SiiD      ------NTGGTIQRYYKVIIAFDVN----EDDLRWLKPGMTVDASVITGKHSIMEYLLSPLMK----GVDK------------AFSEPVNTKRLDTP----------------------------------------------------
LipC      ------KANGEPYYQMQVTVSPEGMKMLSG---EDIKPGMPVEVFVKTGSRSLLSYLFKPILD----RAHT------------SLTEE-------------------------------------------------------------
EmrA      ALVPRDVSAGEFTK---VVQRIPVKIKITKGDLSLLRVGMGGEVEIRRTR---------------------------------------------------------------------------------------------------
Spr0693   SP-AAGNNTGSKYPY-TIDVTG---------EVGDLKQGFS--VNIEVKSKT--KAILVPVSSLVMDDSKNY--VWIVDEQQKAKKVEVSLGNADAENQEITSGLTNGAKVISNPTSSLEEGKEVKADEATN-----------------
MacA      -----EKVNDAIFYYARFEVPN---------PNGLLRLDMTAQVHIQLTDVK--NVLTIPLSA-LGDPVGDNRYKVKLLRNGETREREVTIGARNDTDEIVKGLEAGDEVVIGEAK---PGAAQ-------------------------
AcrA      SDVTVDQTTGSITLR--AIFPN---------PDHTLLPGMFVRARLEEGLNP--NAILVPQQGVTRTPRGDATVLVV-GADDKVETRPIVASQAIGDKWLVTEGLKAGDRVVISGLQKVRPGVQVKAQEVTADNNQQAASGAQPEQSKS
HlyD      ------QKLGLVFN---VIVSVEENDLSTGNKHIPLSSGMAVTAEIKTGMRSVISYLLSPLEE----SVTE------------SLHER-------------------------------------------------------------
Cons.     ------43553262---64533---------0012+3459421829435200--112102000----0001------------10100-------------------------------------------------------------
```

**S. Fig. 2.**

**S. Fig. 2.** Sequence alignment of SiiD with six structurally homologous PAPs (UniProt accession no. in parentheses): LipC (Q54457), EmrA (O67159), Spr0693 (A0A0Y3EE53), MacA (P75830), AcrA (P0AE06) and HlyD (P09986). Helical (H in red), extended β-structure (E in blue) and other (-) types of secondary structure predicted for SiiD are shown above the SiiD sequence. Predicted transmembrane helices are highlighted in red. Domain boundaries of LipC, EmrA, Spr0693, MacA, AcrA and HlyD were derived from their 3D structures and highlighted in light blue (β-barrel domain), in dark blue (lipoyl domain) and in green (coiled-coil domain). Amino acid residues at the turning in the individual coiled-coil domains are highlighted in magenta (in the Spr0693 structure some residues (underlined) are missing in this region). The HlyD structure was solved without the β-barrel domain. Highlighted in yellow are the N- and C-terminal residues of the three investigated variants SiiD(30-393), SiiD(43-300) and SiiD(80-263). The conservation score (Cons.) displays the conservation of physicochemical properties for each column of the alignment (values from 0 to 10, 10 is represented by "+", "*" means 100% conservation, and columns labeled with "-" exhibit > 25% gaps).

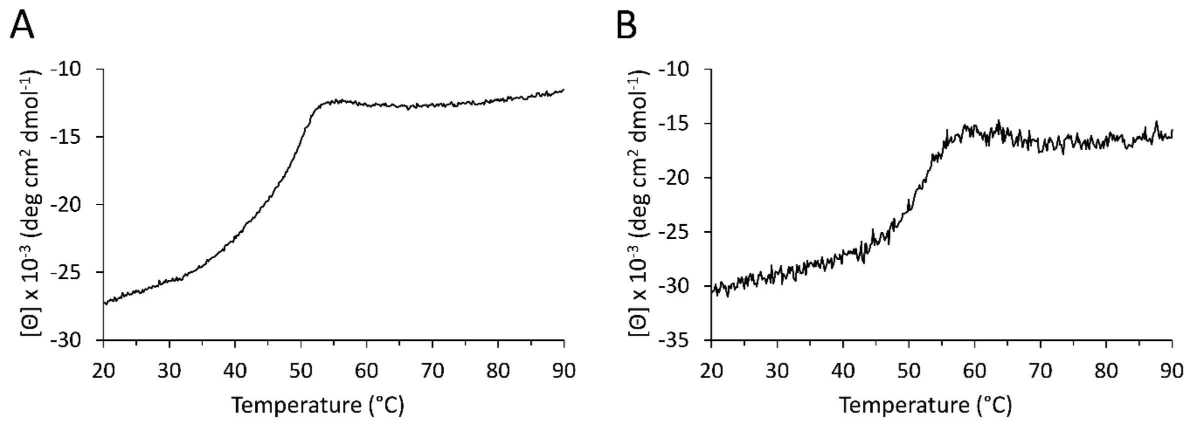


**S. Fig. 3.** Curves of ellipticity as a function of temperature for variants (A) SiiD(43-300) and (B) SiiD(80-263) recorded at 222 nm and 220 nm, respectively.

```
          1       10        20        30        40        50        60        70
                     HHHHHHHHHHH        HHHHHHHH          HHHHHHHHHH   EEEEE
SiiF      MDKKLEPYYLSAETALSIVSKKFNIKIDIKEDDINLRFKKYDRNNTDDSIQMKNFFLSLGLSLQDILFNN
PCAT1     ----QYDLTDCGAACLSSIAQYYGLKMSLA--KIREMTGTD-TQGT-NAYGLIHAAKQLGFSAKGVKA-S
                     HHHHHHHHHH   B   HH  HHHHHH          HHHHHHHHHHH    EEEE
                  20        30        40           50         60        70

                   80            90        100       110       120          130
                     EEEEE       EEEEEEEE    EEEEE GGG  EEE HHHH        EEEEE
SiiF      GEDL-LNEPMPILLLTPEM----KWMVCVSG-GQKIKLVNARGELCYVEIEDEYLKELS---AFSILPL
PCAT1     KEDLLKDFRLPAIANVIVDNRLAHFVVIYSIKNRIITVADPGKG--IVRYSMDDFCSIWTGGLVLLEPG
                     EEEE         EEEEEEEE  EEEEE        B   HHHHHH    EEEE
             80        90       100       110         120       130       140

                    140       150       160       170       180             688
                         HHHHHHH HHHHHHHHHHHHHHHHHHHHGGGHHHHHHH
SiiF      NKVVDS---IRVKNIIKNSLSMNKIFYTKYFFSSLFMAIFALTIPVFSNLFYDK...............
PCAT1     EAFQKGDYTQNMMVKFAGFLKPLKKTVLCIFLASLLYTALGIAGSFYIKFLFDD...............
               B     HHHHHHH  GGGHHHHHHHHHHHHHHHHHHHHHH HHHHHHH
               150       160       170       180       190                  727
```

**S. Fig. 4.** Sequence alignment (excerpt) of SiiF and PCAT1 (UniProt accession no. A3DCU1) produced by SWISS-MODEL. Shown are the N-terminal domain (approximately up to residue 130 in SiiF) and the first transmembrane α-helix (underlined in PCAT1) of the transmembrane helix domain. Secondary structure elements for the SiiF model and the PCAT1 3D structure were calculated using the DSSP program (displayed are α-helices (H in red), $3_{10}$-helices (G in green), extended β-structure (E in blue), and β-bridge (B in blue)). Highlighted in yellow are the N- and C-terminal residues of the three investigated variants SiiF(2-120), SiiF(2-124) and SiiF(2-128). Highlighted in orange is the catalytic essential cysteine of C39 peptidases (Cys21 in PCAT1).

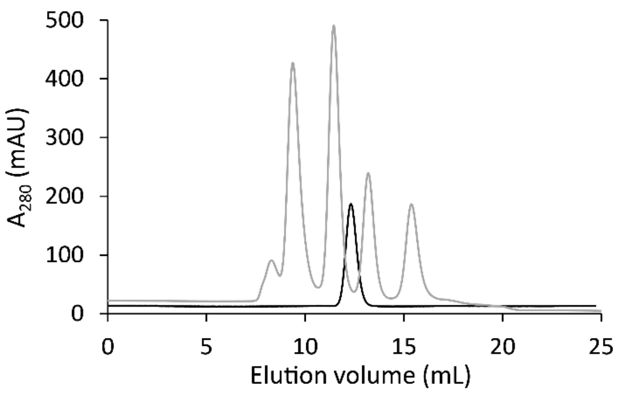


**S. Fig. 5.** Analytical size exclusion chromatography of SiiF(2-128). The absorbance spectrum of the sample is shown together with the elution profile of molecular weight standards (grey curve, major peaks from left to right: conalbumin (75 kDa), carbonic anhydrase (29 kDa), ribonuclease A (13.7 kDa) and aprotinin (6.5 kDa)).

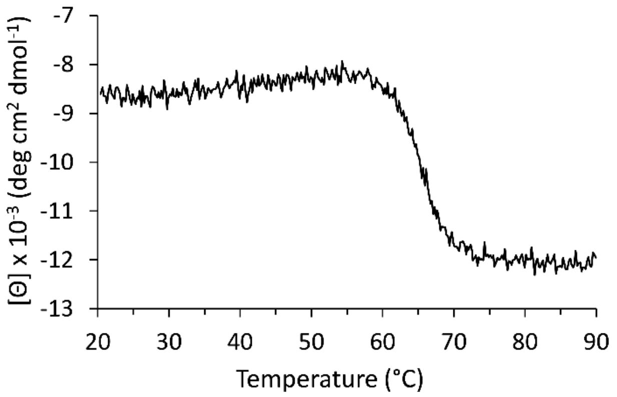


**S. Fig. 6.** Curve of ellipticity as a function of temperature for variant SiiF(2-128) recorded at 215 nm.
